# Cerebrospinal Fluid and Interstitial Fluid Motion via the Glymphatic Pathway Modelled by Optimal Mass Transport

**DOI:** 10.1101/043281

**Authors:** Vadim Ratner, Yi Gao, Hedok Lee, Maikan Nedergaard, Helene Benveniste, Allen Tannenbaum

## Abstract

It was recently shown that the brain-wide cerebrospinal fluid (CSF) and interstitial fluid exchange system designated the ‘glymphatic pathway’ plays a key role in removing waste products from the brain, similarly to the lymphatic system in other body organs^1,2^. It is therefore important to study the flow patterns of glymphatic transport through the live brain in order to better understand its functionality in normal and pathological states. Unlike blood, the CSF does not flow rapidly through a network of dedicated vessels, but rather through peri-vascular channels and brain parenchyma in a slower time-domain, and thus conventional fMRI or other blood-flow sensitive MRI sequences do not provide much useful information about the desired flow patterns. We have accordingly analyzed a series of MRI images, taken at different times, of the brain of a live rat, which was injected with a paramagnetic tracer into the CSF via the lumbar intrathecal space of the spine. Our goal is twofold: (a) find glymphatic (tracer) flow directions in the live rodent brain; and (b) provide a model of a (healthy) brain that will allow the prediction of tracer concentrations given initial conditions. We model the liquid flow through the brain by the diffusion equation. We then use the Optimal Mass Transfer (OMT) approach^3^ to model the glymphatic flow vector field, and estimate the diffusion tensors by analyzing the (changes in the) flow. Simulations show that the resulting model successfully reproduces the dominant features of the experimental data.

## 1. INTRODUCTION

In this paper, we describe a mathematical model of the glymphatic systems using the theory of optimal mass transport. Cerebrospinal fluid (CSF) is continuously produced and transported into and out of the brain via the glymphatic pathway, a process which also facilitates removal of toxic waste proteins^1–3^. Notably, soluble amyloid beta (Aβ) oligomers and tau protein have been shown to clear via the glymphatic pathway^1,4^ and it has been hypothesized therefore that dysfunction of glymphatic transport underlie buildup of extracellular Aβ aggregates and tau as seen in Alzheimer’s disease. The anatomical structure of the glymphatic pathway is complex but a key component is the perivascular compartment. The outer perimeter of the perivascular space is defined by glial endfeet with high expression of aquaporin 4 channels that facilitate convectively driven CSF movement into the interstitial fluid (ISF) space which result in clearance of ISF waste solutes via peri-venous conduits (Figure 1). Glymphatic transport of fluid and solutes is driven by vascular pulsations and most importantly by a pressure gradient over the cerebral convexity (so-called ‘transmantle’ pressure gradient) which can be manipulated, even by minor physiological alterations such as shifting body posture^4^.

**Figure 1:**
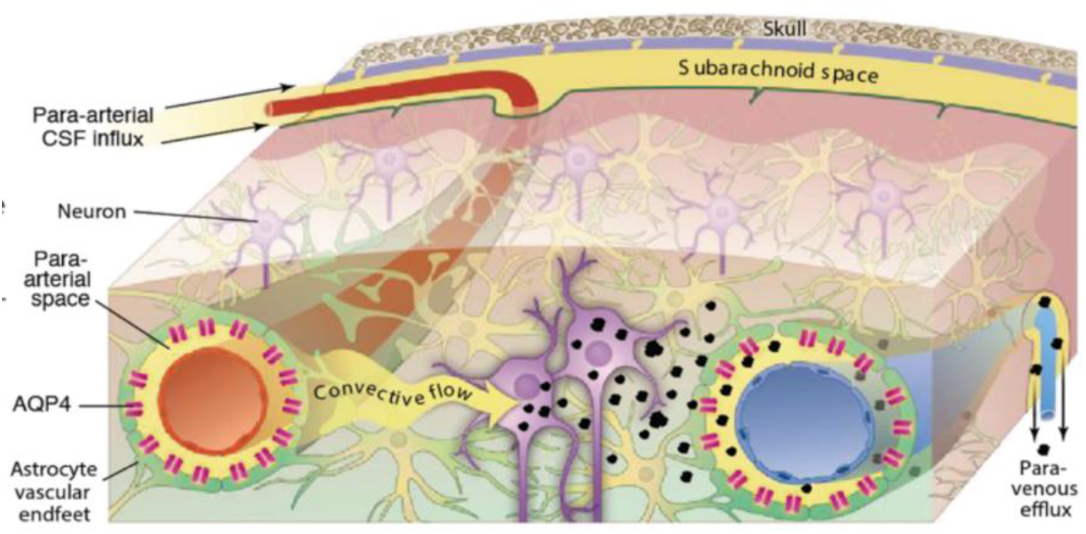
Schematic of the glymphatic pathway in rodent brain. Para-arterial inflow of cerebrospinal fluid (CSF) enters the brain tissue faciliated by astrocytic AQP4 water channels. The CSF mixes with the interstitial fluid (ISF) and propels the waste products of neuronal metabolism into the paravenous space, from which they are directed into cervical lymphatic vessels and ultimately returned to the general circulation for clearance by the liver.

Initial studies of the glymphatic pathway used fluorescently tagged optical dyes of different molecular weight (MW) administered into CSF in combination with in vivo two-photon microscopy.This technique allowed for dynamic characterization of the fast peri-arterial influx of tracers dyes in small cortical areas of rodents, and also defined the MW cut off for transport of solutes from the peri-arterial space and into brain parenchyma (ref). Most importantly, intra-parenchymal injections of radiolabeled or fluorescently tagged tracers in combination with optical imaging showed that soluble Aβ cleared along the peri-vascular space of large central veins.

The limitation of optical *in vivo* imaging in regards to understanding the brain-wide glymphatic pathway is the relatively small areas of the brain which can be studied. This lead to supplementary optical *ex vivo* studies of sectioned brain slices analyzed at varying times after CSF administration of the dyes (ref). These studies demonstrated that small MW tracers were able to distribute within the entire perivascular glymphatic pathway of the mouse brain within 30-60 min. However, quantification of glymphatic waste *clearance* from the live brain including visualization of this dynamic ‘macroscopic’ process on a system level was difficult to assess using the optical imaging techniques. This lead to development of magnetic resonance imaging (MRI) techniques.

In the present work, we derive mathematical models of the glymphatic system based on an optical flow model derived from optimal mass transport ideas, by which one can construct a continuous transition from two time points while preserving “mass” (e.g., image intensity) during the transition. The theory even allows a short extrapolation in time. This technique will be explicated in Section 2 below. Preliminary results on this topic have been reported in some previous work^12^.

## 2. BACKGROUND ON GLYMPHATIC PATHWAY

The first MRI study used paramagnetic contrast (Gd-DTPA, MW 938 Da) administered into the CSF of a rat in combination with contrast-enhanced, dynamic T1-weighted MRI. A series of T1 weighted 3D MRIs of the rat brain visualized in real time how the contrast molecule moved into the entire rat brain along the large arteries at the base and that contrast slowly (over 2-4 hrs) penetrated into the brain parenchyma. Quantification of glymphatic transport of Gd-DTPA was accomplished using both parametric ROI analysis as well as non-parametric cluster analysis. Another recent study extended this analysis to include 2-compartment modeling where ‘retention’ and clearance of a given tracer was calculated over a finite period of time in the brain and in pre-selected brain regions such as the cerebellum or the hippocampus. Although these analytical approaches were useful to compare global measures of glymphatic transport between rats and across groups they were not able to spatially or dynamically map the glymphatic transport process.

More specifically, in the central nervous system (CNS), clearance and/or transport of protein and excess solutes from the interstitial fluid (ISF) space is less well understood but is thought to be governed in part by cerebrospinal fluid (CSF) exchanging with ISF via para-arterial conduits (Virkow-Robin spaces)^1^. In recent studies using optical imaging in combination with fluorescently tagged tracers injected into the intrathecal space, this CSF-ISF exchange system was further characterized in the live rodent brain^2^. Specifically, it was demonstrated that small molecular weight (MW) tracers injected into the subarachnoid CSF move rapidly into the brain along both penetrating cortical and basal arteries (but not veins) to reach the capillary beds and associated interstitial compartments ^2^ (Figure 1). It has been confirmed that the fluorescently tagged tracers administered into CSF are cleared from the brain along the same pathways as intra-parenchymally injected tracers, accumulating primarily along large draining veins. It was also discovered that astrocytes play a pivotal role in linking the para-arterial inflow and the para-venous efflux pathways^2^. In mice harboring the deletion of the astroglial aquaporin 4 (AQP4) water channels, tracer influx along para-arterial routes and through the brain interstitium was sharply reduced suggesting that fluid is mass transported through the astrocytic network. Together, the paravascular influx and clearance routes, coupled by trans-astroglial fluid mass transport, constitute a brain-wide waste clearance pathway, which is designated the ‘glymphatic system’^2^. To translate the glymphatic pathway findings observed with fluorescently tagged tracers and optical imaging towards a relevant preclinical testbed, intrathecal injection of paramagnetic contrast molecules was combined with dynamic T1-weighted 3D brain MRI on a 9.4TmicroMRI instrument in rodents.

In the rodent brain, the primary, most rapid glymphatic inflow (defined as total normalized uptake >100) occurs at the level of the hypothalamus, olfactory, retrosplenial cortex, pons, amygdala, cerebellum and the hippocampus. Slower influx and clearance of Gd-DTPA is observed in the entorhinal cortex, insula and more remote brain regions such as caudate putamen, midbrain and thalamus. Importantly, the contrast-enhanced MRI technique for quantifying glymphatic pathway function was validated using conventional ex vivo optical imaging and fluorescently tagged tracers of same molecular weight^11^. These studies show that we can quantify and capture both influx and efflux components of glymphatic transport the live rodent brain using contrast-enhanced MRI^11^.

Driven by the initial image data of flow in the glymphatic system, we propose the use of optimal mass transport to try to estimate the velocity vector field, and in turn employ this methodology this to derive better computational fluid models, especially modelling the diffusion and advection fluxes at various point in the brain. Having a mathematical model based on the underlying physics of fluid flow, would give us a better biological understanding of what occurs in the glymphatic systems, and will allow us to propose further experiments, as well as predict how changes in the parameters (e.g., cell volume, pulsatility, AQP4 channels, and supine/prone positions) effect the overall flow.

We will sketch the mathematics of optimal mass transport in the next section, use this to derive an analytic model of the transport through the *glymphatic system* based on fluid flow physics and brain physiology, and then test and validate our model on a dynamic series of contrast-enhanced MRI images in Section 4.

## 2. OPTIMAL MASS TRANSPORT (OMT)

### 2.1 Assumptions about the Data

The data is collected as a time series of T1-weighted MRI images taken after administration of paramagnetic contrast (Gd-DTPA, molecular weight 938 Da) into the lumbar intrathecal space of an anesthetized rat. The dynamic acquisition of data depicts a gradual propagation of the tracer into the brain. Based on anatomical mapping of signal changes induced by Gd-DTPA on the MRI images over time we assume (***assumption 1***) that the tracer is transported via the glymphatic pathway. We further assume (***assumption 2***) that brain-wide glymphatic transport can be described by the diffusion equation with anisotropic tensor coefficients that do not change with time (or change very slowly). The problem of flow estimation and modeling is then translated into the inverse problem of estimating the diffusion coefficients from concentration data. The model estimation process consists of two steps: an optical flow computation and diffusion tensor estimation; each of which are described below.

We finally assume (***assumption 3***) that the total mass of the tracer agent in the brain is almost constant (mass preservation) between consecutive time points. This assumption was made in order to simplify the optical flow estimation (and because modeling the tracer inflow and decay is not an easy task). A new model that takes tracer mass changes into account is currently being developed.

### 2.2 Optical Flow

At first, optical flow is computed between each pair of consecutive MRI images^4,5^. Optical flow may be defined as the apparent motion of brightness patterns in a sequence of intensity images. In this work, we have employed an optical flow model based on optimal mass transport (OMT) also known as the Monge-Kantorovich problem^3,6^. It employs an energy minimization principle with a mass conservation constraint, and thus is physically very natural.

More specifically, let Ω_0_ and Ω_1_ denote two subdomains of **R**^*d*^, having smooth boundaries, each with a positive density function, *μ*_0_ and *μ*_1_, respectively. We assume ∫_Ω_0__ *μ*_0_ = ∫_Ω_1__ *μ*_1_ = 1 so that the same total mass is associated with Ω_0_ and Ω_1_ (assumption 3). We consider diffeomorphisms *ϕ*: (Ω_0_, *μ*_0_) → (Ω_1_, *μ*_1_), which map one density to the other in the sense that

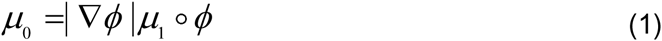

(Jacobian equation). A mapping *ϕ* that satisfies the Jacobian equation is said to have the *mass preservation* (MP) property, written as *ϕ* ∈ MP. Here |∇*ϕ*| denotes the determinant of the Jacobian map ∇*ϕ*. A mapping *ϕ*. that satisfies this property may be considered as a redistribution of the mass of material from one distribution *μ*_0_ to another distribution *μ*_1_. There are in general many such mappings, and we want to pick out an optimal one. Accordingly, the optimization in the *L*^*p*^ Kantorovich – Wasserstein metric

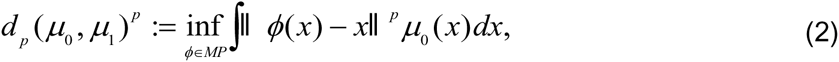

seeks an *optimal* MP map which, when it exists, minimizes a cost functional. This functional is seen to place a penalty on the distance the map *ϕ* moves each bit of material, weighted by the material's mass. The cases *p* = 1,2 have been extensively studied. For *p* = 2, a fundamental result is that there is a unique optimal *ϕ* ∈ MP transporting *μ*_0_ to *μ*_1_, and that this *ϕ* is the gradient of a convex function *u*, i.e., *ϕ* = ∇*u*. Thus, the Kantorovich-Wasserstein metric defines as distance between two mass densities the cheapest way to transport one mass distribution to the other. The optimal transport map, denoted by *ϕ*_*MK*_. When *p* = 2, the optimal transport map may be computed by solving a PDE^3^.

## 3. ANALYTICAL MODEL OF THE BRAIN

Let *μ*(*t*, **x**) be a nonnegative function describing the density of the tracer over the 3D brain domain, in time. We assume that tracer propagation through the brain tissue can be described by the following diffusion equation (assumption 2):

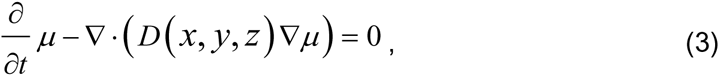

where ∇. and ∇ denote divergence and gradient with respect to spatial coordinates, respectively, and *D* is a (spatially varying) symmetric 3 × 3 matrix (diffusion tensor). The eigenvectors of *D* point in the main diffusion directions, with the eigenvalues denoting the diffusivity (diffusion strength) in those directions. Our goal is to find *D*, given the transfer speeds defined for each MRI time sample by the mass transport maps *ϕ*_*MK*_. Once *D* is known, we can compute tracer concentration at any point in time and space given any initial (and boundary) conditions. Given a single transport map, we can only find the strength, *k*, of diffusion in the direction of the transport vector, *w*, in each voxel

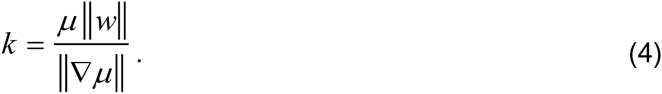

This single direction-value combination does not provide sufficient information about the diffusion tensor. An analysis of all such combinations, however, reveals a more complete picture. At first, we construct a set Σ for each pixel, composed of vectors whose directions are those of the transport vectors (*w*) originating from that pixel at different times, and their magnitudes – the corresponding scalar diffusivities (*k*). For every *v*∈Σ, we also add –*v*∈Σ, since the diffusion direction is unsigned (i.e. diffusivity in one direction is the same as that in the opposite direction). The principal components of Σ are therefore parallel to the eigenvectors of the (approximated) diffusion tensor *D*, and their variances are the corresponding eigenvalues.

We will next show the application of these ideas to the modelling of the glymphatic pathway of a rodent brain.

## 4. EXPERIMENTAL RESULTS

Four rats were used for the analysis given below. We give now the pre-processing descriptions and then the results comparing the conventional and optimal mass transport (OMT) results. As we will see these results while consistent with one another also give complementary information.

### 4.1 Brief Processing Description

The dynamic T1-weighted MRI time series for each of the rat was processed as follows. Briefly, general MRI image processing consisted of head motion correction, intensity normalization, smoothing and voxel-by-voxel conversion to ‘% of baseline signal’ using SPM8. Using PMOD software the dynamic time series was converted in 4D data and overlaid on the corresponding T1 weighted anatomical MRI. A 3D mask was generated based on the anatomical template and used to extract the brain-relevant information from the dynamic 4D time series. The same anatomical mask was used to generate the OMT data for each rat. 3D visualization of contrast distribution in the rat brains was executed using Amira software. ‘Conventional’ contrast distribution was displayed against the OMT based visualization of fluid dynamics in the whole brain at corresponding time points after contrast administration into the cisterna magna. The two analyses from four different rat brains are displayed below in Figure 2.

**Figure 2:**
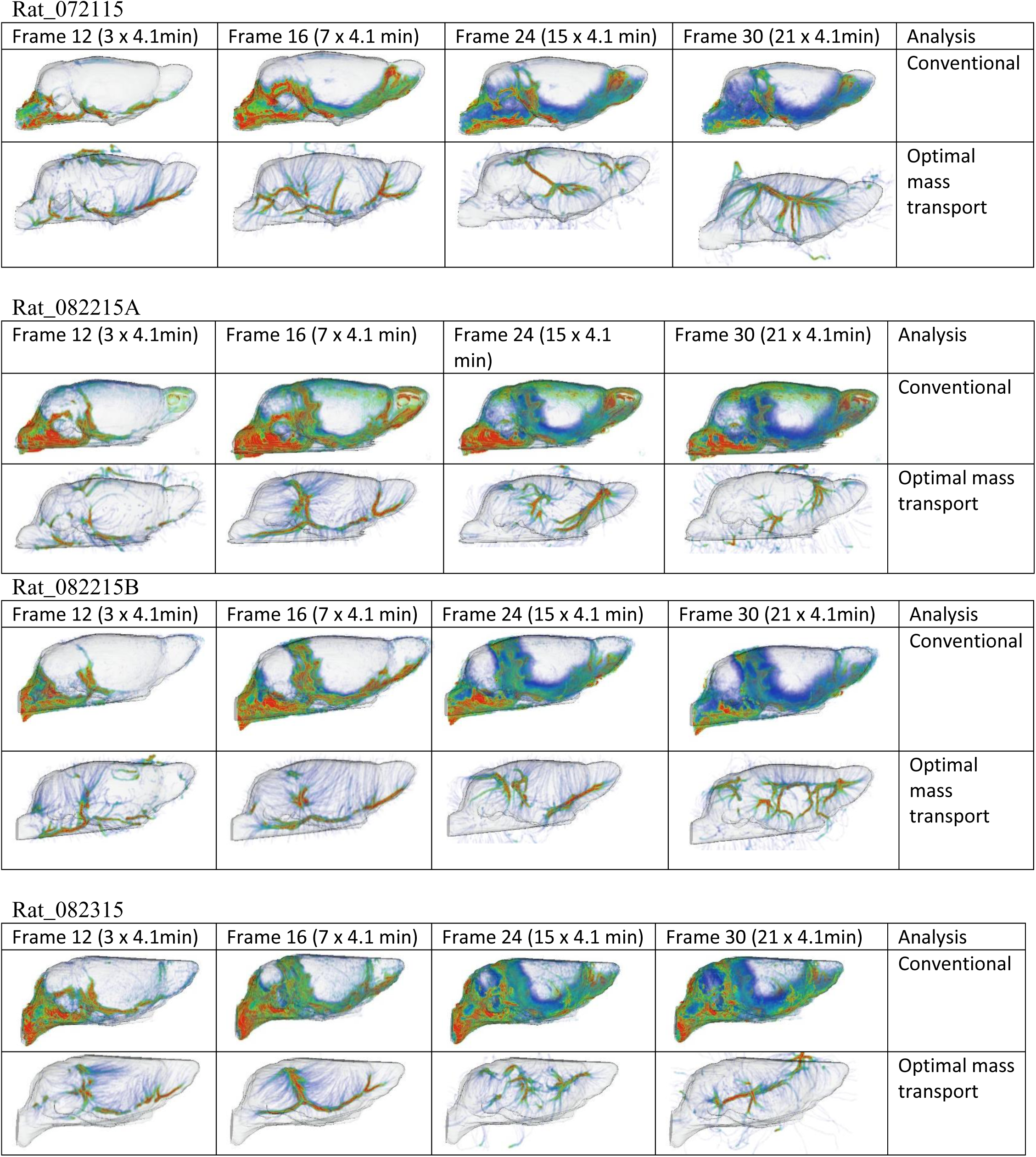
Comparison of conventional and OMT based analyses of the glymphatic pathway in a rat brain.

### 4.2 Description of Results

#### Conventional contrast analysis

the conventional contrast distribution patterns over time are very similar from rat to rat. At early times after Gd-DTPA administration into CSF via the cisterna magna, contrast is most prevalent in areas associated with the large arteries at the base of the brain as previously described (ref). For anatomical orientation, the color coded map of contrast distribution is displayed on a volume rendered surface mask of the whole rat brain (red and blue colors represent high and low contrast, respectively). Over time, contrast moves into the brain parenchyma via the glymphatic pathway. The parenchymal contrast uptake is evident as a slow penetrating ‘gradient’ moving from basal to more central and olfactory areas of the rat brain.

#### OMT based analysis

The OMT analysis provides supplementary information on ‘mass’ (contrast) transfer when compared to the conventional analysis. At early time points, the OMT ‘pathways’ (displayed in color code) shows strong mass transfer along the base of the brain in areas associated with the large arteries similar to the conventional analysis. However, parenchymal ‘streams’ of mass transfer from the base of the brain towards the surface of the brain and towards the olfactory bulb is evident on the OMT processed brain but not on the conventional analysis. At later time points the OMT analysis is further informative by showing strong stream lines in brain areas associated with large veins. This is very promising given that mass / solute clearance from the brain is occurring along peri-venous spaces (ref). Although the OMT analysis is slightly different between the individual rats all show the same trend i.e. at early times (first 20-40 min) strong mass transfer along large arteries at the base of the brain and in the fissure between the cerebellum and forebrain where large arteries are also present. At later times the clearance process become more evident and the OMT provides anatomical information on where the clearance occurs, i.e., in central areas along large venous complexes.

## 5. DISCUSSION

A number of other researchers have modeled liquid propagation by diffusion equations including those in hydrogeology and brain research^7,8^. Most of these studies attempt to solve the estimation problem for a domain with constant, or locally constant coefficients. In the present work, we present the first attempt (to our knowledge) to fully solve the inverse problem of dense estimation of coefficients that are either anisotropic or spatially-varying or both, from concentration measurements. Also, none of the previous attempts to solve the inverse problem (assuming simpler, homogeneous models) utilized OMT.

OMT provides a good estimate of the actual glymphatic flow. Despite its simplicity, the proposed model mimics the real data closely, with some errors arising from the difficulty of the inverse problem. Some refinement of the diffusion model, such as including additional terms in the diffusion equation and compensating for the mass source (and loss of mass) while computing the OMT flow, should improve its accuracy.

## ACKNOWLEDEMENTS

This research was partially supported by the National Center for Research Resources under Grant P41-RR-013218, the National Institute of Biomedical Imaging and Bioengineering under Grant P41-EB-015902 of the National Institutes of Health through the Neuroanalysis Center of Brigham and Women's Hospital, National Institutes of Health grant 1U24CA18092401A1 through Stony Brook Medical School, and the Air Force Office of Scientific Research though grants FA9550-09-1-0172 and FA9550-15-1-0045.

